# The titin N2A-MARP signalosome constrains muscle longitudinal hypertrophy in response to stretch

**DOI:** 10.1101/2025.06.19.660595

**Authors:** Robbert van der Pijl, Jochen Gohlke, Joshua Strom, Eva Peters, Shengyi Shen, Stefan Conijn, Zaynab Hourani, Stephan Lange, Ju Chen, Paul Langlais, Siegfried Labeit, Henk Granzier, Coen Ottenheijm

## Abstract

Titin-based mechanosensing is a key driver of trophic signaling in muscle, yet the downstream pathways linking titin sensing to muscle remodeling remain poorly understood. To investigate these signaling mechanisms, we utilized unilateral diaphragm denervation (UDD), an in vivo model that induces titin-stiffness-dependent hypertrophy via mechanical stretch. Using UDD in rats and mice, we characterized the longitudinal hypertrophic response and distinguished stretch-induced signaling from denervation effects by performing global transcriptomic and proteomic analyses following UDD and bilateral diaphragm denervation (BDD) in rats. Our findings identified upregulation of titin-associated muscle ankyrin repeat proteins (MARPs). Subsequent phosphorylation enrichment mass spectrometry in mouse diaphragm highlighted the involvement of the N2A-element. UDD in MARP knockout (KO) mice resulted in enhanced longitudinal hypertrophy, with Western blot analysis revealing activation of the mTOR pathway. Furthermore, pharmacological inhibition of mTORC1 with rapamycin suppressed longitudinal hypertrophy, demonstrating that mTOR signaling regulates titin-mediated hypertrophic growth in a MARP-dependent manner. These findings establish MARPs as key modulators of titin-based mechanotransduction and highlight mTORC1 as a central regulator of longitudinal muscle hypertrophy.

## INTRODUCTION

Titin is a giant protein in striated muscle and forms an elastic filament in sarcomeres^1^, the smallest contractile unit in muscle. One of the functions attributed to titin is to generate passive tension in muscle^2,3^, ensuring optimal overlap of the actin-based thin filaments and myosin-based thick filaments for muscle contraction. The elastic properties of titin combined with its arrangement in the sarcomeres, spanning the half-sarcomere, make titin an ideal stress sensor. The mechanosensory properties of titin have been extensively described^4–6^ and appear to localize to several signaling hotspots on titin.

In skeletal muscle, the N2A element is the prevalent titin region associated with hypertrophy signaling. The N2A element is known for its interaction with the muscle ankyrin repeat proteins (MARP1-3)^7,8^ and calpain3^9^. All MARPs have been implicated in trophicity signaling, with redundancy between the MARP proteins^7,10^. MARP1 tethers titin to the thin filament, forming a mechanism for increasing passive tension^11,12^. In the heart, MARP1 also interacts with MLP and protein kinase C alpha in intercalated disks, resulting in a maladaptive trophic response in dilated cardiomyopathy^10^. MARP2 can be phosphorylated at serine-99 (S69 in mice) by Akt, resulting in reduced differentiation potential of myoblasts^13^. MARP2 has also been linked to the NFkB-pathway in inflammatory responses^14^, suggesting MARP2 may have roles in atrophy signaling. MARP3 is the least studied member of the MARPs but appears to play a role in glucose uptake and vascular remodeling in skeletal muscle^15,16^. Titin has previously been shown to activate signaling in response to stretch^17,18^, providing a potential activation mechanism for N2A to activate trophic signaling.

Muscle hypertrophy is directly associated with titin-based stiffness. High stiffness, as observed in the Ttn^Δex112–158^ and Ttn^Δex219-225^ mouse models^19,20^, results in longitudinal hypertrophy, i.e. an increase in serial-linked sarcomeres, to reduce sarcomere length and normalize passive tension. We previously used a surgical model in which we denervated one hemi-diaphragm (unilateral diaphragm denervation, UDD)^21^, inducing passive cyclic stretch of the denervated hemi-diaphragm by the innervated costal to induce (longitudinal) hypertrophy. This hypertrophy was dependent on titin-based stiffness, as higher titin stiffness resulted in exaggerated hypertrophy and lower titin stiffness resulted in attenuated hypertrophy^21^. UDD is a unique model as it allows the *in vivo* study of passive stretch-induced muscle hypertrophy.

We used UDD as a model for studying titin-mechanosensing and examined the signaling that underlies the hypertrophy response. Our findings support the N2A-MARP signalosome inhibiting longitudinal hypertrophy and show mTOR signaling to be a prominent contributor to longitudinal hypertrophy.

## RESULTS

### Longitudinal hypertrophy regulates transient trophicity in UDD

A unique aspect of unilateral diaphragm denervation (UDD) is the transient nature of the hypertrophy. This transient nature is likely the result from changes in fractional extension of sarcomeres. Sarcomere length (SL) at end-expiration length was 2.9±0.1 µm in sham, while stretch lengths at end-inspiration were predicted to reach 3.7 µm directly after denervation of the right costal diaphragm (FIG. 1A, previously published^21^). Such sarcomere lengths put high strain on titin (FIG. 1A, right panel), providing a potent trigger for mechanosensing and subsequent hypertrophy signaling. Our results show that during 6-days of UDD the denervated costal rapidly hypertrophies, followed by slow onset of atrophy (FIG. 1B). The mass increase coincides with longitudinal hypertrophy of muscle fibers, addition of serial-linked sarcomeres, adding 952±81 sarcomeres (30.2% increase in total fiber length; p<0.0001; FIG. 1C) following 6-days of UDD. This increase in sarcomere number appeared to stabilize by 6-days UDD, as at 12-day UDD sarcomere addition was measured to be 778±87 sarcomeres versus sham levels (not significantly different compared to 6-day UDD). Assuming that fractional extension of sarcomeres during inspiration decreases with longitudinal hypertrophy, 30.2% increase in sarcomere number would reduce the sarcomere length at end inspiration from 3.7 µm to ∼2.8 µm, in close agreement with whole body formaldehyde perfusion experiments which showed sarcomere lengths of 2.7±0.1 µm^21^. As a SL of 2.8 µm falls below the end-expiration SL, the denervated costal effectively no longer experiences “stretch”, reducing titin’s mechanosensing trigger and thus explaining the transient nature of the hypertrophy seen in UDD.

**FIGURE 1.**
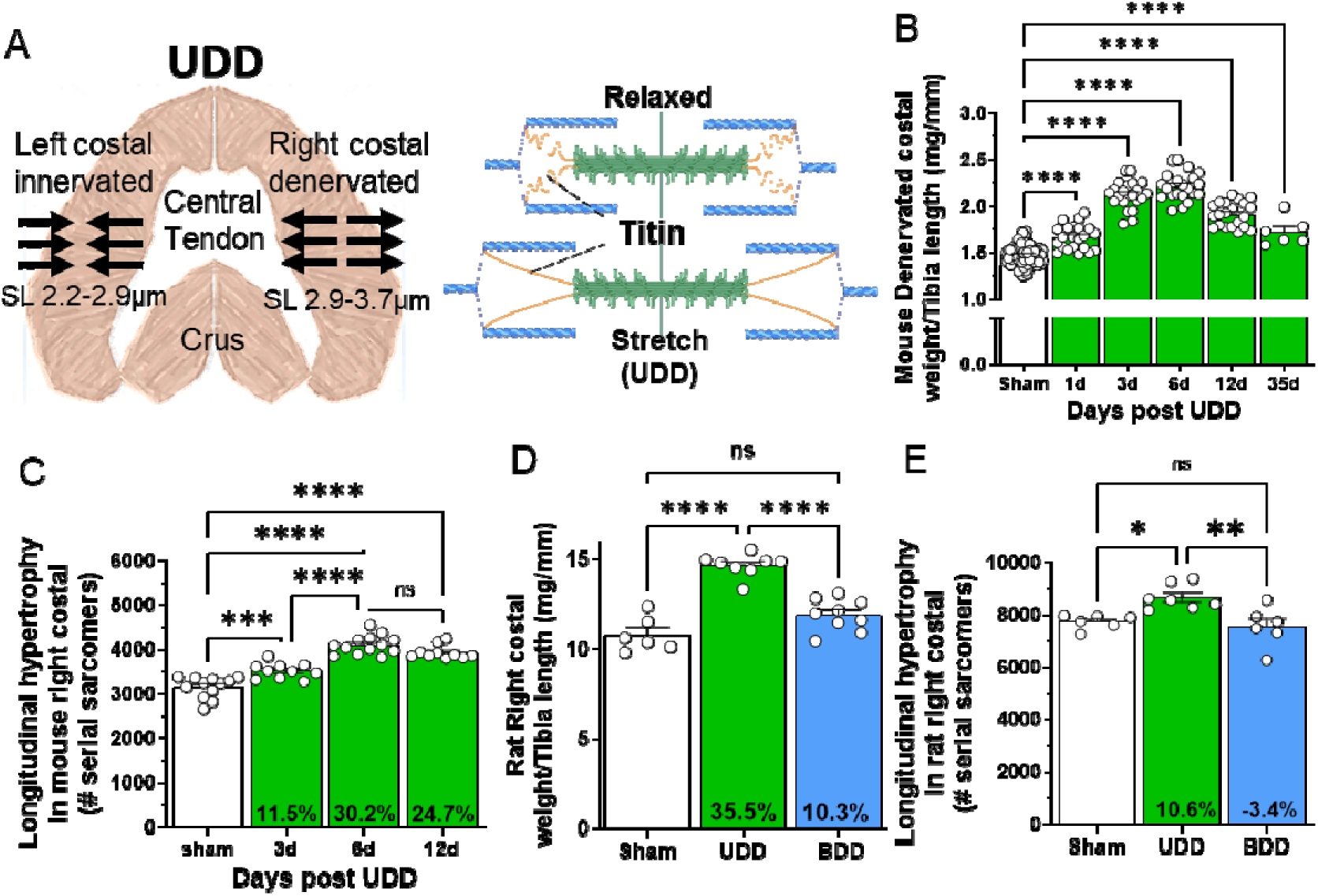
Transient hypertrophy following UDD in the denervated costal diaphragm. (A) schematic of how UDD affects sarcomere length in the denervated costal (left panel) and how stretch extends titin for mechanosensing (right panel, generated through BioRender). (B) Mouse diaphragm costal weight normalized to tibial length following 1, 3, 6, 12 and 35-days of UDD, showing the hypertrophy phase peaking at 6-days and progressing to the atrophy phase at 12-days post-UDD (n=6-22, shams grouped for simplicity). A substantial part of the hypertrophy seen in mice encompasses longitudinal hypertrophy (C), lengthening of the muscle fibers by addition of serial sarcomeres (n=10-13). The increased fiber length likely reduces the stretch-based hypertrophy signaling and thus explains the transient nature of hypertrophy. (D) 3-day BDD in rats (n=6-9) confirms stretch is the trigger for inducing hypertrophy in UDD at the tissue mass level (D; rat diaphragm right costal normalized to tibial length, denervated in UDD) and serial sarcomeres level (E). One-way ANOVA, with Tukey post-hoc testing.

To confirm that stretch is the trigger for hypertrophy, rather than denervation, we performed bilateral diaphragm denervation in rats (BDD), resulting in complete inactivity of the diaphragm. Note that we attempted BDD in mice but experienced high mortality rates, while survival in rat was >75%. All rats were of similar body weight pre-surgery and showed a slight decrease in body weight after 3-day BDD (Supplemental FIG. 1A; p=0.036), and comparable tibial length and soleus weights (Supplemental FIG. 1B & C). Similar to mice, rats present with increased denervated costal mass after 3-day UDD: 14.68±0.64 mg/mm versus sham rats 10.82±0.39 mg/mm (FIG. 1D; p>0.0001; muscle weight normalized to tibial length). However, 3-day BDD rats did not develop increased mass 11.92±0.89 mg/mm compared to sham rats 10.82±0.39 mg/mm. The increase in costal mass at 3-day UDD results in part from an increase in longitudinal hypertrophy, with denervated costal diaphragm adding 933±301 serial sarcomeres compared to sham animals (FIG. 1E; p=0.018), whereas 3-day BDD rats showed no difference in serial sarcomere number (7559±313 versus sham 7740±111). Thus, stretch and not denervation triggers longitudinal hypertrophy.

To support titin’s role as mechanosensor of stretch to drive hypertrophy, we performed RNAseq on both UDD and BDD operated rat diaphragms. Splicing appears mostly unaffected in UDD animals, while BDD rats showed marked increased splicing in exons encoding the elastic PEVK region of titin suggesting reduced titin compliance (Supplemental FIG. 2). To study how titin compliance might affect the hypertrophy response in UDD, we performed UDD on Rbm20^ΔRRM^ mice and Rbm20 ko rats. Rbm20 is an RNA splicing-factor for titin and both the mouse and rat model generate larger, more compliant titin isoforms and should thus be less sensitive to stretch. Rbm20^ΔRRM^ mice showed relatively less tissue mass increase compared to wildtype (WT) 3, 6, or 12-days UDD (Supplemental FIG. 3A), without changes in longitudinal hypertrophy at 6-days UDD (Supplemental FIG. 3B) suggesting the difference in mass is a result from radial fiber growth, in agreement with our previous findings^21^. Finally, we compared denervated costal mass from Sprague Dawley (SD) and Rbm20 ko rats 3-day UDD, showing a difference in mass increase of 24.4±3.8% in Rbm20 ko and 35.5±5.9% in SD rats (p=0.02; Supplemental FIG. 3C), supporting that stretch evokes hypertrophy in muscle and that titin compliance modulates the extent of hypertrophy across species. Thus, muscle stretch in the costal diaphragm is a potent trigger for longitudinal muscle hypertrophy, with titin being a primary sensor.

**FIGURE 2.**
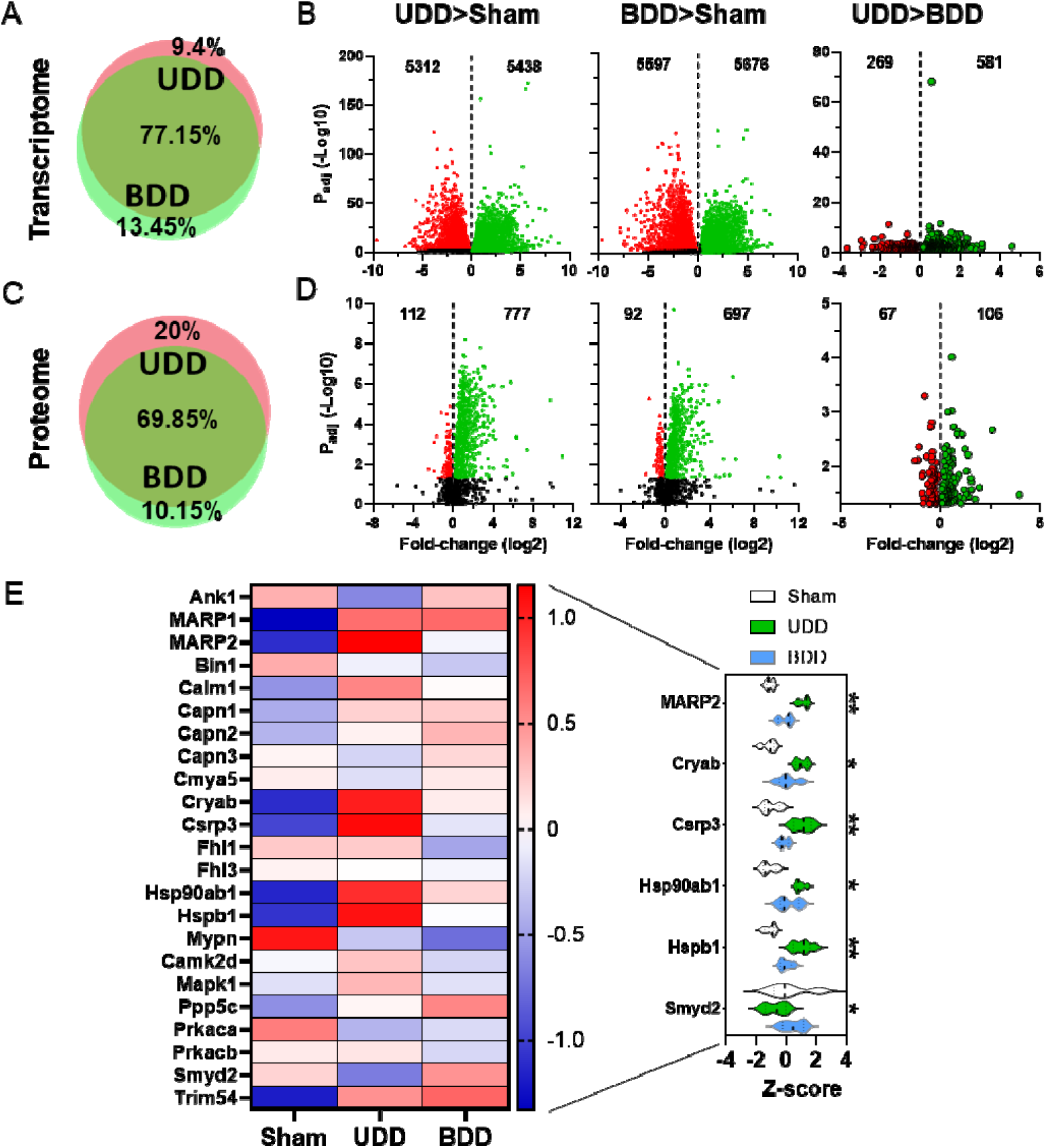
Global transcriptomics and proteomics following 3-days UDD and BDD in rats. Global transcript studies by RNAseq of sham, UDD and BDD right costal diaphragm (n=5/group). Same parameters apply to the global proteome studies (C-D) with mass spectrometry. Quantitative Venn diagrams of the transcriptome (A) showing overlap gene regulation between UDD and BDD. Vulcano plots of UDD (B, left) and BDD (B, middle) showed similar gene regulation. Comparing UDD to BDD directly revealed just 850 differentiall regulated genes (B, right) indicating a small subset being responsible for hypertrophy regulation. Quantitative Venn diagrams of the proteome (C) showed similar regulation compared to transcriptome. Volcano plots of UDD (D, left) and BDD (D, middle) showed primarily upregulation of proteins. Comparing UDD to BDD directly revealed just 173 differentiall regulated proteins (D, right). Green-dots: upregulated genes/proteins, red-dots: downregulated genes/proteins. Titin-associated proteins in heatmap of proteome (E; Z-score: red= upregulated, blue= downregulated) and violin plots (right panel) of differential proteins between UDD (red) and BDD (blue), indicating upregulation of titin-associate proteins following stretch. 2-way ANOVA (sham vs UDD and sham vs BDD) p_int_: *p<0.05, **P<0.01.

**FIGURE 3.**
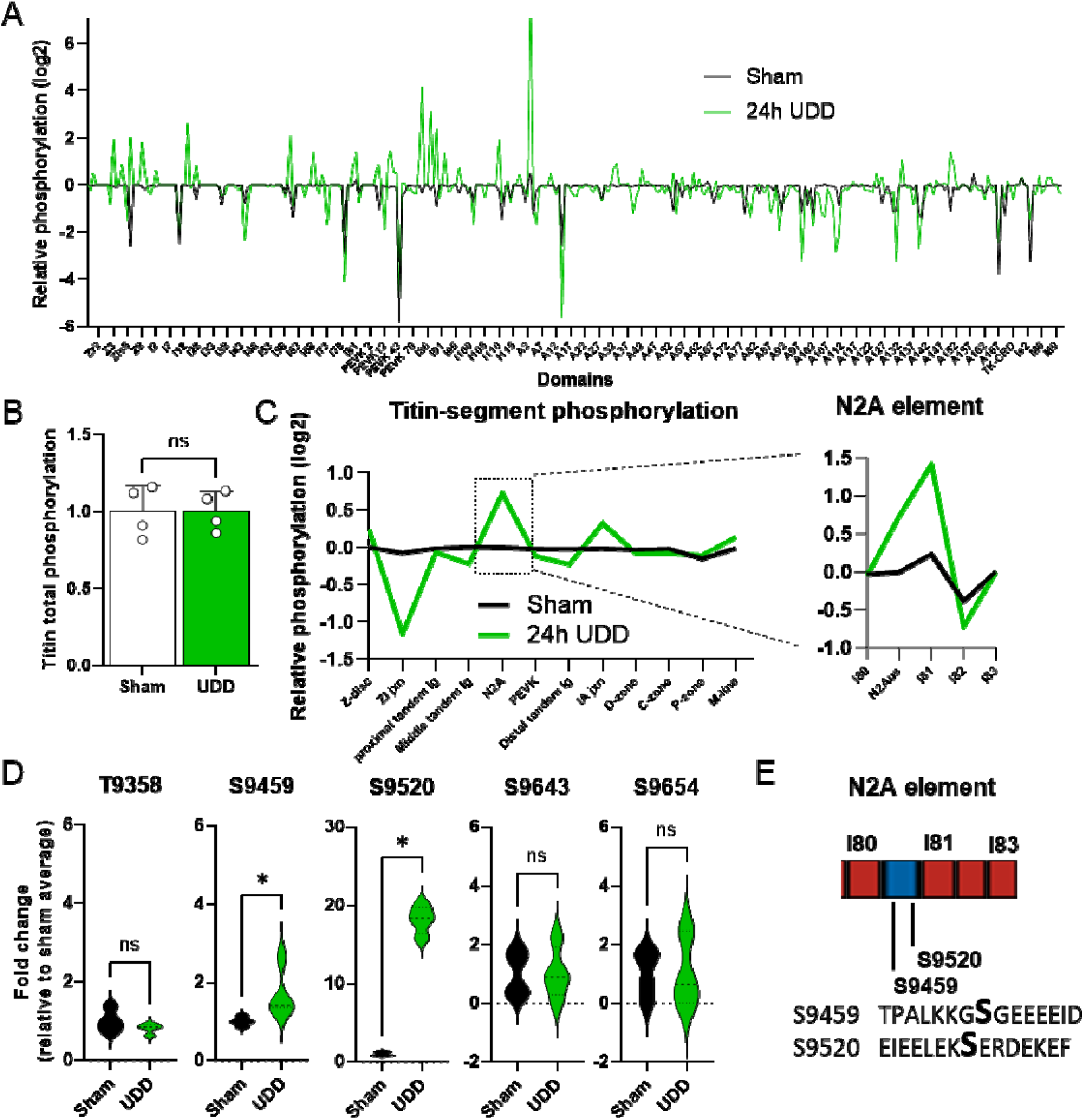
Phosphorylation of titin following 24-hours of UDD by mass spectrometry (n=4/group). Phosphorylation of individual titin domains (A) relative Z-score (log2) of titin phosphorylation showing domain specific changes in phosphorylation. Total phosphorylation of titin (B) is not affected by UDD, however titin showed regional changes in phosphorylation (C), notably increased phosphorylation of the N2A-element (boxed). Fold-change of the phosphorylation signal for the 5 main sites found in the N2A-element (D). (E) Schematic of the N2A element with the 2 pSer found in the N2Aus (Transcript: ENSMUST00000099981.10 Ttn-203). Red: Ig domain coding and blue: unique sequence coding. Mouse titin phosphorylation and global mass spectrometry was analyzed by t-test, Kolmogorov-Smirnov test or multiple t-test with a cut-off at P<0.05.

### Global transcriptome and proteome level studies reveal a distinct subset of genes and proteins involved in UDD stretch hypertrophy

To identify the cellular processes involved in stretch-based hypertrophy we performed both transcriptome wide studies by RNA sequencing (RNAseq) as well as proteome studies by global mass spectrometry (MS) on our rat 3-day sham, UDD and BDD samples. Initial studies focused on separating the effect of denervation versus stretch. Principal component analysis of the top 500 genes showed close clustering of samples by group (Supplemental FIG. 4A), suggesting distinct gene programs are active between sham, UDD and BDD. At the protein level, principal component analysis of the top 250 proteins suggests narrow separation of groups, predicting overlap of the BDD and UDD samples (Supplemental FIG. 4B). Gene expression studies revealed very close overlap of BDD and UDD profiles with 77.15% overlap in differentially expressed genes (DEG’s: p_adj_<0.05) and just 9.4% of DEG’s being specific for UDD (FIG. 2A). Volcano plots of UDD>Sham (FIG. 2B, left panel) and BDD>Sham (FIG. 2B; middle panel) showed the ∼11.000 DEG’s (Supplemental data Tables 1-6) to be nearly equally distributed between up and downregulated DEG’s. Direct comparison of UDD>BDD (FIG. 2B, right panel) revealed 850 DEG’s to be uniquely associated with UDD, of which 581 were upregulated (Supplemental data Table 3). MS studies broadly agreed with the RNAseq data, showing UDD and BDD share overlapping expression programs (FIG. 2A & C). Twenty percent of the differentially expressed proteins (DEP’s; P<0.05) detected are unique to UDD (FIG. 2C). Volcano plots of UDD>Sham (FIG. 2D, left panel) and BDD>Sham (FIG. 2D, middle panel) showed 889 and 789 DEP’s, respectively, (Supplemental data Tables 7-12) which were primarily increased. Comparing UDD>BDD revealed 173 DEP’s (FIG. 2D, right panel) that are unique to UDD. Most of DEP’s (and DEG’s) found are related to transcription, translation, energetics and folding, with the highest fold DEP’s in rat UDD>BDD being: Med15 (15.5-fold), Eif3i (6.0-fold) and Gtpbp3 (3.9-fold) (Supplemental data Table 9). GOterm enrichment of UDD>BDD, separated by downregulated and upregulated DEG’s (Supplemental FIG. 3C, left top and bottom graph, respectively), and DEP’s (Supplemental FIG. 4C, right top and bottom graph, respectively), indicated that upregulated DEG/DEP’s are primarily related to muscle development and function consistent with an active hypertrophy program and downregulated targets with metabolism and cellular respiration. Concurrently, KEGG PATHWAY database searches (Supplemental data Table 6) indicated that the DEG’s are involved in muscle remodeling. Thus, UDD has a expression profile that is distinct from denervation (BDD), with more focus on remodeling of muscle.

Exploring if titin-mechanosensing was active in either UDD or BDD we generated heatmaps of titin-binding proteins based on RNAseq (Supplemental FIG. 5) and global MS data (FIG. 2E). To narrow down proteins that were differential between UDD and BDD we tested (2-way ANOVA p_int_<0.1) the z-score and found: MARP2 (Ankrd2; p=0.003), Cryab (p= 0.026), Csrp3 (p= 0.002), Hsp90ab1 (p= 0.09), Hspb1 (p= 0.008) and Smyd2 (p= 0.004), to be unique to UDD. Interestingly, these DEP’s (MARP2, Smyd2, Hsp1b, Hsp90ab and Cryab) are established binding partners of the N2A element of titin.^22^

### Titin phosphorylation in UDD

With the rat data indicating changes in titin-associated signaling and titin stiffness modifying the hypertrophy response in mice, we evaluated how UDD affects titin post-translationally. We performed MS on phosphorylation enriched peptides from 24-hour UDD diaphragm samples. 24-hour UDD was selected to study the early phosphorylation events. MS revealed 2870 phosphorylated peptides (p<0.05), of which 142-sites were in titin (∼700-sites identified including non-significant sites; Supplemental data Table 13). We opted to define titin phosphorylation by analyzing the phosphorylation at the domain level to study if titin showed regional changes in phosphorylation (FIG. 3A and Supplemental data Table 14), as total titin phosphorylation (FIG. 3B) was unchanged. Domain level analysis indicated titin phosphorylation was primarily changing in the I-band. This motivated quantifying titin phosphorylation by segment (FIG. 3C and Supplemental data Table 15; following established naming conventions in Kolmerer and Labeit^1^, and Bang et al^23^). Segmental analysis of titin (FIG. 3C) revealed a marked increase in phosphorylation of the N2A-element and I/A-junction, and a decrease in the Z/I-junction. Focusing on the increase in phosphorylation of the N2A-element, a known trophicity signaling hub (reviewed in ^22^), we assessed the specific domain phosphorylation of the N2A-element (FIG. 3C, highlight). In the N2A-element we found 7 phosphorylation sites (FIG. 3D-E) of which S9346 & S9350 are located in a linker sequence between I79 and I80, S9483 in I80, S9459 & S9520 in the N2A unique sequence, S9654 in I81 and S9643 in I82). The 2 sites located in the N2A unique sequence were significantly upregulated following UDD (FIG. 3D-E). pS9459 and pS9520 currently have no known function but could serve as a recruitment signal (see discussion).

### The MARP proteins regulate hypertrophy in UDD

The MARP1 and MARP2 proteins were strongly expressed both at the RNA level and protein level following UDD (FIG. 2E). MARPs have been shown to be important regulators of trophicity and have been proposed as intermediaries between titin and trophic signaling pathways (see discussion). As the MARPs share high homology and possible redundancy^7^, we performed 6-day UDD on both single- and multi-KO combinations of the MARPs. We initially performed UDD on the MARP triple KO mice (MARP tKO; knockout of Ankrd1, -2 and -23 genes) and found a 12% reduction in the hypertrophy response compared to WT (FIG. 4A; p=0.0099). This reduction in hypertrophy supports a potential link between titin-mechanosensing and MARP-based trophic signaling. To test if one or a combination of MARPs caused the reduction in hypertrophy, we performed 6-day UDD on MARP1 (Ankrd1), MARP2 (Ankrd2) and MARP3 (Ankrd23) KO mice (FIG. 4B-D) and double KO for MARP1/2, MARP1/3 and MARP2/3 (Supplemental FIG. 6). Whereas MARP1 KO mice did not show a difference in denervated costal diaphragm mass, MARP2 KO mice showed a 14% increase (p=0.003) in mass and MARP3 KO mice showed a 13% reduction (p=0.004) in mass gain. However, all MARP double KO mice also presented with attenuated hypertrophy (Supplemental FIG. 6). This suggests that the separate MARP proteins could have distinct functions in UDD, and there may be a dependency for two MARPs in regulating trophic signaling, further discussed in the discussion section.

**FIGURE 4.**
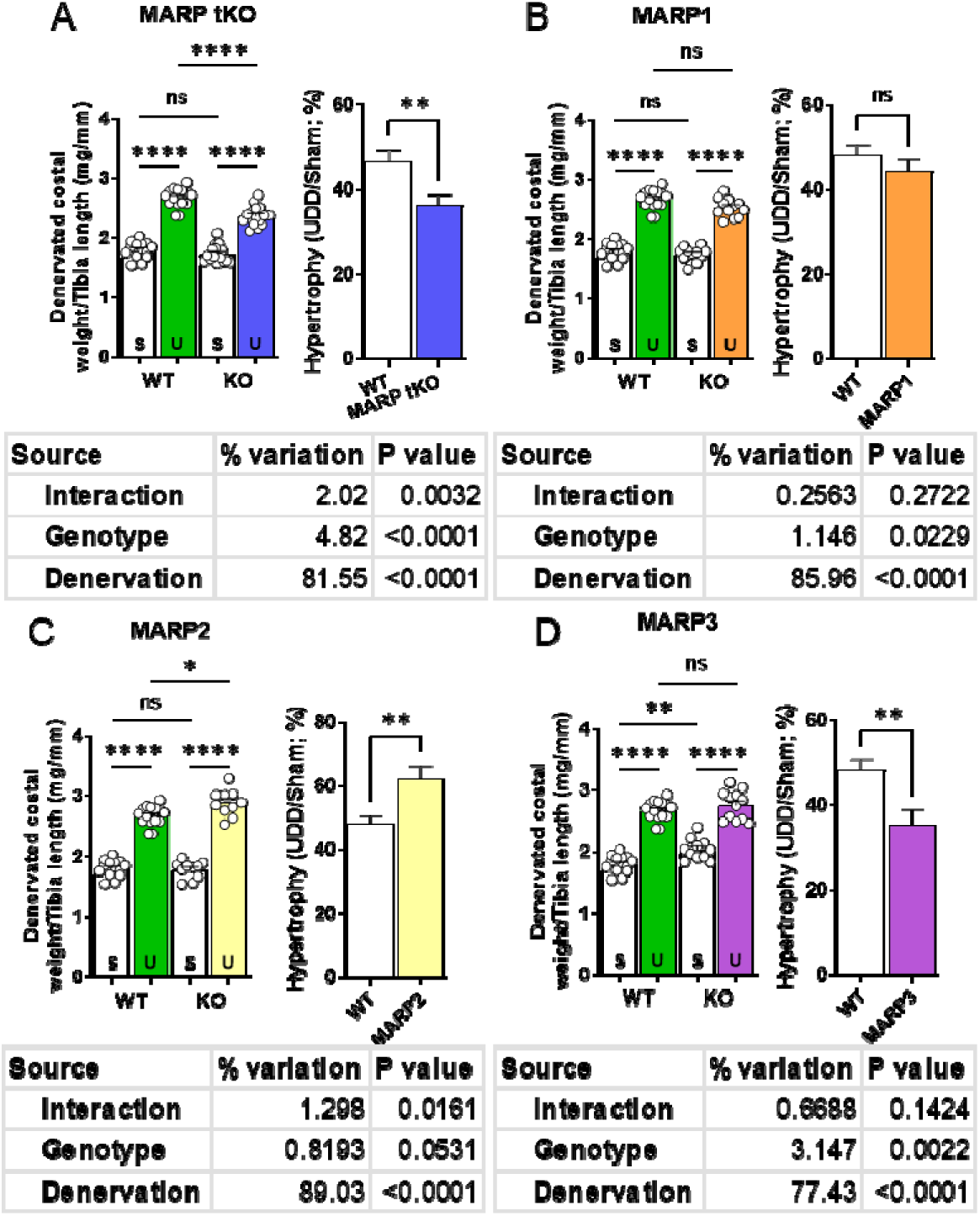
6-day UDD on KO mice of MARP proteins. MARP triple KOs show a reduced response to UDD (A; p<0.01). No effect of single MARP1 KO (B) on UDD, increased hypertrophy following MARP2 KO (B; p<0.01; t-test), indicating possible roles in hypertrophy suppression or atrophy signaling and MARP3 KO (C) showed baseline hypertrophy in costal diaphragm in addition to less hypertrophy development in UDD (p<0.01; t-test) compared to WT, implying roles as a suppressor of hypertrophy. Left panel, diaphragm right costal mass normalized to tibial length and right panel, percentual increase in right costal mass relative to sham. S= Sham, U= UDD (n=10-12). Statistical testing by t-test or two-way ANOVA with Tukey post-hoc testing.

### MARPs negatively regulate longitudinal hypertrophy

Prompted by the reduction in diaphragm mass gain during UDD in MARP tKO mice and by the complexity of targeting all the single MARP KOs, we assessed longitudinal hypertrophy in 6-day UDD MARP tKO samples. MARP tKO mice at baseline have fewer serial sarcomeres than WT (2476±55 vs. 3157±69, respectively; p<0.0001), however 6-days UDD MARP tKO had similar numbers of sarcomeres to WT, 4081±44 versus 4109±59 (FIG. 5A). These data show MARP tKO mice add 653 ± 91.6 more sarcomeres than wildtype mice (FIG. 5B; 1605±71 vs. 952±59, respectively; p<0.0001). To discern a possible mechanism of the MARPs inhibiting longitudinal hypertrophy, we probed several candidate proteins of hypertrophy pathways that were prominent in the MS datasets. Western blots for MAPK1/3, Calcineurin, mTOR, P70 S6K and 4E-BP1 were performed on costal diaphragms of WT and MARP tKO, 6-day Sham and UDD animals. All samples were normalized to Gapdh and data were presented as relative to WT sham (FIG. 5C; representative western blot images in FIG. 5D). Mapk1 showed increases in protein level that were comparable between WT and MARP tKO, whereas MAPK3 showed preferential upregulation in MARP tKO (p= 0.0099). Calcineurin was unchanged following UDD, note that in a t-test the WT mice showed a significant increase in UDD. In WT mice, mTor showed a striking increase in expression following 6-day UDD (p= 0.0006), whereas in MARP tKO this was trending (P= 0.08). Downstream proteins of mTORC1; P70 S6K and 4E-BP1 both showed altered regulation, with P70 S6K being similarly upregulated in WT (p= 0.0005) versus MARP tKO (p= 0.001), while 4E-bp1 was only significantly upregulated in MARP tKO (p= 0.033). 2way-ANOVA for the signaling proteins did not indicate specific pathway changes between WT and MARP tKO.

**FIGURE 5.**
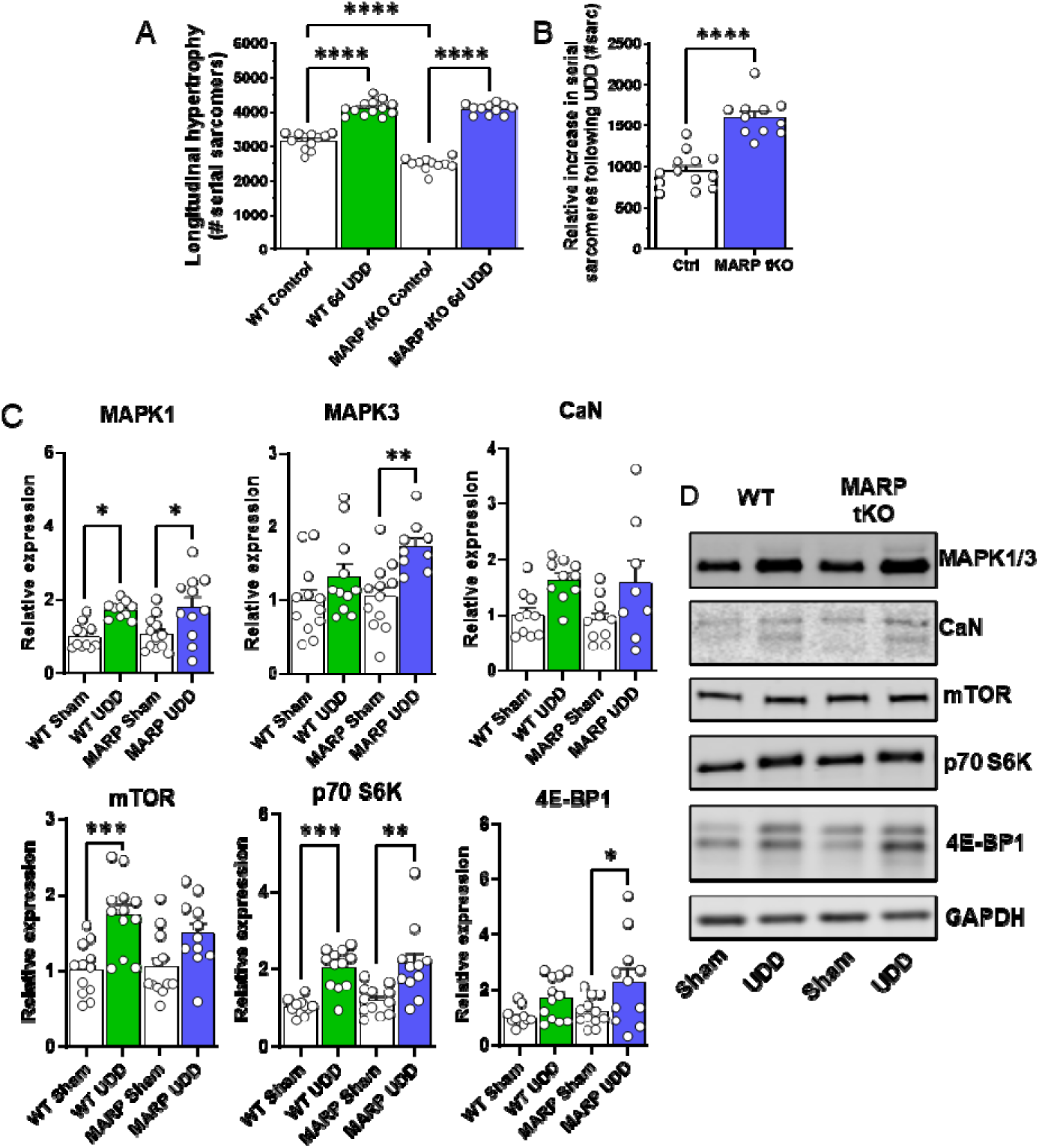
MARPs inhibits longitudinal hypertrophy. Longitudinal hypertrophy measured in right costal strips of WT and MARP tKO mice in 6-day sham and UDD mice (A). Numerical increase in serial sarcomeres is higher in MARP tKO (P<0.0001) mice compared to WT mice (B; n=7-12), suggesting that the MARPs inhibits longitudinal growth. Probing hypertrophy signaling by western blot, normalized to Gapdh, with expression set relative to WT sham levels (C; n=8-12). Differential mTor response suggests role in regulating longitudinal hypertrophy. (D) Representative blot images of the signaling proteins. Statistical testing by one-way or two-way ANOVA with Tukey post-hoc testing.

### mTOR signaling regulates longitudinal hypertrophy in UDD

Motivated by the data from the mTor western blots in the MARP tKO mice and mTor being a prominent regulator of skeletal muscle hypertrophy, we performed UDD in WT mice treated with rapamycin, an inhibitor of mTORC1 signaling and skeletal muscle hypertrophy. The rat transcriptome studies also indicated calcium signaling (Supplemental data Table 6) to be a prominent pathway in UDD. To test the role of calcium signaling we included a second inhibitor; cyclosporin A, an inhibitor of the calcineurin-NFAT pathway, another hypertrophy regulating pathway. Mice received twice daily intraperitoneal injections of either rapamycin (2.5 mg/kg/day), cyclosporin A (25 mg/kg/day) or vehicle (DMSO), starting 3-days prior to surgery until sacrifice of the mice (schematic in FIG. 6A). Cyclosporin A treated mice did not show a change in hypertrophy (FIG. 6B) or serial sarcomere addition (FIG. 6C), indicating that the calcineurin-NFAT pathway is not a primary mechanism for hypertrophy in UDD. Treatment did not affect the innervated left costal diaphragm (FIG. 6D), body weight (FIG. 6E) or tibia length (FIG. 6F). Interestingly, mice that received rapamycin displayed less hypertrophy of the denervated costal diaphragm (FIG. 6B; - 11.2%; p<0.001) compared to vehicle treated mice. This attenuated hypertrophy response coincided with a reduction in serial sarcomere addition (FIG. 5C; - 8.5%; p<0.001) compared to vehicle. Note that following 3-day UDD, rapamycin treated mice did not show significant increases in serial sarcomeres compared to untreated sham animals (Δ67 ± 81 sarcomeres, versus vehicle treated mice Δ363 ± 79 sarcomeres). This suggested rapamycin treatment almost completely inhibited longitudinal hypertrophy and that mTORC1 signaling plays a vital role in longitudinal hypertrophy development. We thus propose that mTORC1 signaling positively regulates longitudinal hypertrophy in skeletal muscle and that titin’s N2A-element tunes the extent of longitudinal hypertrophy through the MARPs to prevent excess hypertrophy (graphic summary in FIG. 6G).

**FIGURE 6.**
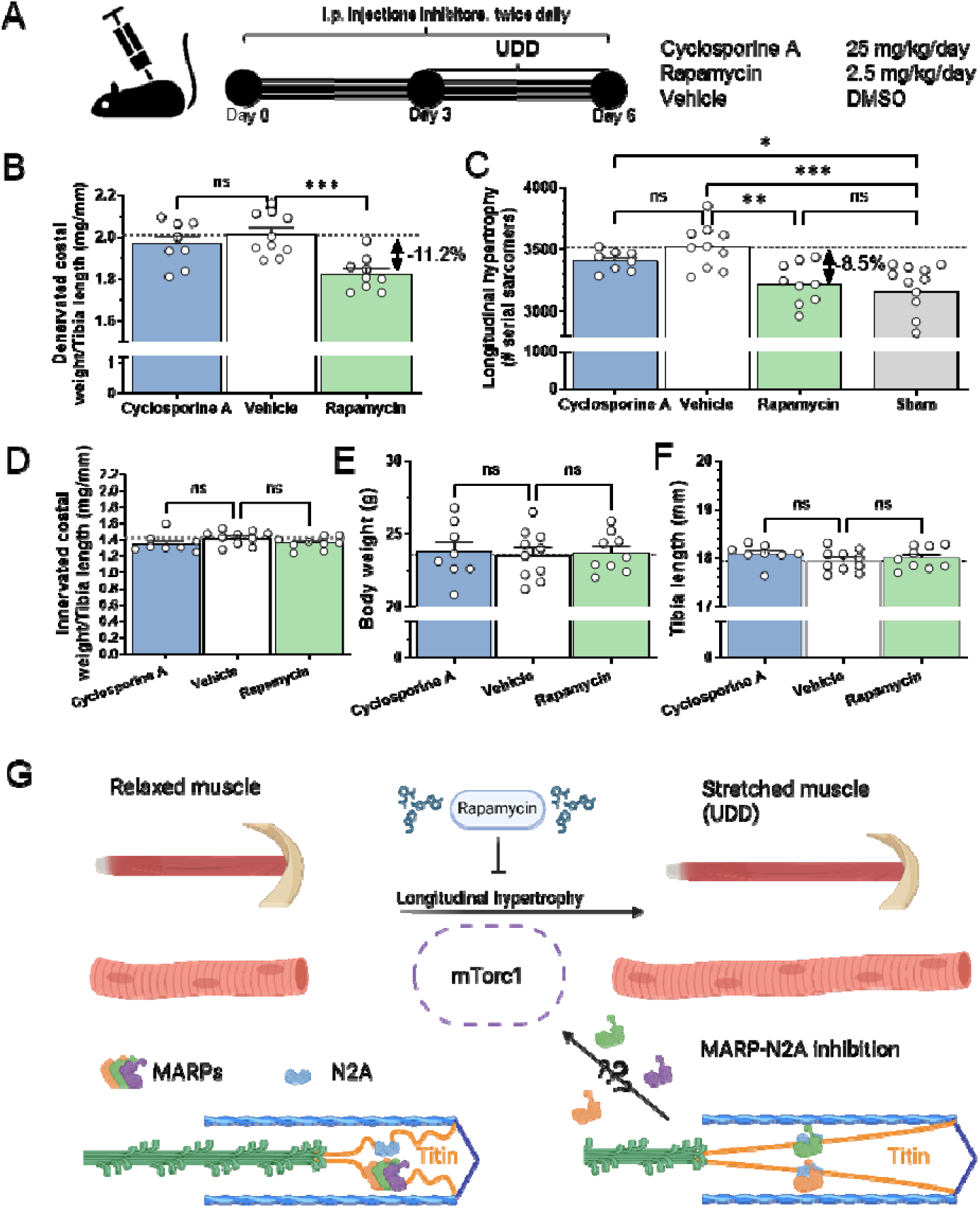
Pharmacological inhibition of the mTor (rapamycin) and calcium (cyclosporin A) based hypertrophy pathways revealed mTor to be involved in longitudinal hypertrophy. (A) Schematic of the inhibition protocol, showing mice were injected with inhibitors for 3-days prior to receiving UDD surgery, with continued twice daily dosing of inhibitors until sacrifice at day 3 post-UDD. Rapamycin inhibited hypertrophy development both at the costal diaphragm mass level (B; p<0.001) and at the longitudinal hypertrophy level (C; p<0.001), whereas cyclosporine A had no effect. Neither cyclosporine A or rapamycin affected the innervated costal diaphragm (D), or body mass (E) and all mice used were of approximately the same size based on skeletal size, as measured by tibia length (F). (G) Hypothetical mechanism for longitudinal hypertrophy following muscle stretch. The mTorc1 pathway is activated by stretch and initiates longitudinal muscle hypertrophy. MARP proteins sequestered by titin’s N2A element are released upon stretch and tunes the longitudinal hypertrophy, thus preventing excessive longitudinal hypertrophy (Image was generated through BioRender). N=8-10/group, statistical testing by 1-way-ANOVA and Dunnett’s multiple comparisons test.

## DISCUSSION

### Longitudinal hypertrophy as a mechanism for reducing stretch-induced mechanosensing

One of the most striking aspects of the UDD surgical model is the early transient hypertrophy. No other muscle denervation model induces hypertrophy, supporting the notion that stretch, even in inactive muscle, is a potent driver of hypertrophic growth. The extreme nature of the stretch at work in UDD, ∼25% stretch of the denervated costal by the innervated costal at a frequency of 120-230 times a minute (respiration rate)^21^, forms a potent trigger for muscle hypertrophy. Following 6-days UDD the denervated costal diaphragm develops 49.7±10.0% (FIG. 1B) increase in mass, which is primarily caused by addition of 952±81 sarcomeres (FIG. 1C) and to a lesser extent by radial fiber growth^21^. The transient nature of the hypertrophy can be explained by the reduction in sarcomere strain due to the addition of sarcomeres in series (longitudinal hypertrophy), removing the “trigger” that underlies the hypertrophy signaling. This hypothesis fits the data (FIG. 1B-C), where before remodeling UDD costal width (i.e., fiber length) is ∼8.7 mm (3000 sarcomeres x SL 2.9 µm), and when stretched 25%^21^ equals a width of ∼10.8 mm. Following 6-days hypertrophic remodeling (UDD), costal width is ∼10.8 mm (4000 sarcomeres x SL 2.7 µm). This suggests that the costal width increase attenuates the hypertrophy trigger caused by stretching, finally resulting in atrophy (FIG. 1B; 35-day UDD). Denervation itself does not appear to induce hypertrophy of the diaphragm, as 3-day BDD in rat showed no hypertrophy (FIG. 1D-E). This does not exclude BDD-mediated denervation from potentially inhibiting or delaying the hypertrophy response. Denervated skeletal muscles have not been reported to experience (longitudinal) hypertrophy. UDD and BDD are both denervation models and hypertrophy occurs specifically in the denervated costal of UDD operated animals. Stretch is thus the mechanical difference between UDD and BDD and the trigger for hypertrophy signaling. BDD rats do experience abdominal breathing, which could stretch diaphragms sufficiently to induce longitudinal hypertrophy after an extended period of BDD. UDD being a denervation model shows that hypertrophy is not necessarily dependent on a muscle’s ability to contract. This indicates that skeletal muscle may derive its signals for hypertrophy during the muscle relaxation-phase when the muscle is at its longest state. Muscle stretch in human is known to benefit muscle growth and is a vital part of exercise routines (reviewed in ^24,25^). Perhaps muscle antagonism may provide sufficient stretch to induce hypertrophy. As contracting muscles inadvertently stretch their relaxed antagonists, they create an elegant feedback loop that promotes both muscle growth and the maintenance of muscle mass.

### Titin’s response to stretch

Titin’s elastic properties have been extensively described^2,3^. These elastic properties are tied to titin’s role in muscle trophicity, having a direct effect on the extent of muscle hypertrophy^20,21,26,27^. We showed that reducing titin stiffness, using both Rbm20 splice-deficient mice and rats, attenuated the stretch-based hypertrophy (Supplemental FIG. 3). Importantly, in a model with increased titin stiffness, it was shown that titin stiffness has a direct effect on the number of sarcomeres in series^20^, indicating that titin is a key regulator of longitudinal hypertrophy. Titin splicing analysis by RNA sequencing did not indicate changes in splicing in response to UDD (Supplemental FIG. 2). Thus, stretch in rat diaphragm does not alter titin-based stiffness. In contrast, we observed reduced incorporation of exons encoding the PEVK region in BDD rats which likely reflects inactivity promoting increased stiffness of titin. The exact mechanism underlying titin-based regulation of longitudinal hypertrophy is unclear and UDD does not alter titin-stiffness at the splicing level. Thus, we studied phosphorylation as a possible mechanism, focusing on signaling hot-spots in titin (reviewed in^5,22,28^) that could mediate longitudinal hypertrophy. Posttranslational changes in titin, particularly phosphorylation, have been studied mostly in relation to passive tension development^29–31^. Specific PTMs for titin have not been widely studied and only recently has ubiquitination been shown to recruit autophagic receptors to the kinase domain of titin^32,33^. Autophosphorylation of titin kinase at Y170 is considered one of the classic mechanosensing responses in titin resulting in phosphorylation of the autophagic receptor Nbr1 at S115/116, activating autophagy signaling in vitro^34^. We did not observe any signs following UDD that titin kinase was autophosphorylated at Y170. This could be related to phospho-peptide abundance being below the detection limit, or that the titin kinase is inactive^35^. In total we identified ∼700 phosphorylation sites in titin of which 142 were significantly affected by stretch. These sites were distributed along the entire length of titin with several hot spots in the PEVK region, likely involved in stiffness regulation, and several in the Z-disk, which could be related to signaling or structural interactions. The sites in the N2A-element (FIG. 3) were of particular interest as previously it has been suggested that S9540 (S9895 according to diaphragm RNAseq by Brynnel et al^20^) could serve as a recruitment signal for MARP1^36^, a protein that is highly upregulated following UDD (FIG. 2E). MARPs have been shown to quench N2A phosphorylation^36,37^, which make the phosphorylation sites in the N2A segment tantalizing targets for studying titin-MARP binding. If S9459 and S9520 (FIG. 3D-E) play such a role remains to be determined, but such insights could provide future avenues for manipulating titin-MARP interactions. Similarly, we found many titin-associated proteins to show differential phosphorylation in UDD, including 3 sites in the MARPs (Supplemental FIG. 5). How these sites contribute to signaling or recruitment to the N2A-element remains to be seen but provide tantalizing targets for follow-up study.

### The MARP proteins and muscle trophicity

Global transcriptome and proteome studies from 3-day UDD showed multiple titin-associated proteins being upregulated (FIG. 2E). We focused our efforts on the MARP proteins, as they are known interacting partners of titin’s N2A-element and have been shown to be important for hypertrophy regulation in the heart^38,39^. We used UDD on MARP KO mice, focusing on the triple KO for MARP1-3 to account for expected redundancy^7,37^. MARP tKO showed a 12% reduction in hypertrophy following 6-day UDD (FIG. 4A). This suggested stretch-mechanosensing operated through the MARP proteins. In an attempt to isolate a single MARP protein as the main effector, we performed UDD on single MARP knock-out mice (FIG. 4B-D). MARP1 KO did not reveal changes in hypertrophy. However, we previously established that MARP1 localizes to the N2A-segment of titin following UDD^21^. We also independently determined that MARP1 cross-links titin to the thin filament to increase passive tension^40,41^, suggesting MARP1 plays mechanical roles over trophic regulation in skeletal muscle. MARP2 KO developed an exaggerated hypertrophy and lastly MARP3 KO showed an attenuated hypertrophy response to UDD, suggesting MARP2 and 3 play opposing roles in hypertrophy regulation. MARP2 interacts with Akt2^13^, providing a tentative link with the mTOR pathway, discussed below. Additionally, MARP2 interaction with P50-NFκB acting as an analogue for IκB^14^ suggests a possible role in inhibiting NFκB atrophy signaling. The role of MARP3 remains incompletely understood, but prior studies indicate that loss of MARP3 enhances glucose tolerance and insulin sensitivity^15^, and MARP3 has been linked to AMPK signaling. AMPK is a key regulator of metabolic pathways and a well-established inhibitor of hypertrophic signaling, in part through suppression of mTOR activity, and is also responsive to mechanical stimuli ^42,43^. To specifically determine if the MARPs affected longitudinal hypertrophy we measured the number of serial sarcomeres across the width of the costal diaphragm in MARP tKO and found that the tKO mice added 653.2 ± 91.55 more sarcomeres than WT following 6-day UDD (FIG. 5B). This suggests that the MARP proteins inhibit longitudinal hypertrophy. A possible mechanism could be that following stretch the three MARPs bind and compete for the N2A-binding site^7,37^ resulting in translocation of specific MARPs for signaling purposes. The benefit of MARP deletion in cardiomyopathy was shown by Lange et al^10^, where deletion of MARP1/2 ameliorated MLP KO induced DCM. MARP KO has not proven detrimental in mice^44^, making this family an attractive target for therapeutic intervention.

### mTOR and longitudinal hypertrophy

Muscle hypertrophy is regulated through a number of pathways, with the insulin-insulin growth factor (IGF) mediated pathway^45–47^ being the most well understood in skeletal muscle and the calcineurin-NFAT pathway in cardiac muscle^48,49^. With most studies focused on radial (cross-sectional) hypertrophy, we aimed to gain insight into the regulatory mechanism underlying longitudinal hypertrophy, two types of hypertrophy that are not necessarily mutually exclusive. The mTOR signaling pathway was a prime candidate as UDD in MARP tKO mice suggested altered mTOR activity (FIG. 5C-D). This prompted us to test inhibition of mTOR signaling and see if mTOR regulates longitudinal hypertrophy. Using rapamycin^50^, an inhibitor that targets the mTORC1 (protein synthesis regulation) complex, we found a reduction in longitudinal hypertrophy following 3-days UDD compared to vehicle treated mice (FIG. 6B). This strongly suggests that the mTORC1- pathway is in-part responsible for longitudinal hypertrophy following UDD. While rapamycin primarily inhibits mTORC1, rapamycin-insensitive signaling through mTORC2 may also contribute to UDD stretch-induced responses. Denervation itself may activate mTORC2-associated signaling pathways, which could confound the interpretation of stretch-specific versus denervation-mediated mTOR responses in this model. Therefore, a contribution of rapamycin-insensitive mTOR signaling to the observed phenotype cannot be excluded. Although we focused on mTOR signaling in longitudinal hypertrophy development, we do not exclude other pathways being important. mTOR formed an attractive target as previous work showed that longitudinal stretch phosphorylates Akt and upregulates MARP2^51^, forming a tentative link between longitudinal stretch, MARPs and the mTOR pathway. mTORC1-based hypertrophy is primarily mediated through P70 S6K and 4E-BP1 and their respective phosphorylation signals at T389 and T37/46^46,47^. These sites were unchanged at 24-hour UDD (supplemental table 13) indicating that these sites are activated at a later stage, as reported by Norrby et al^52^, or that mTOR is activated through alternative pathways following stretch. Further studies are needed to establish the roles of the various pathways in stretch hypertrophy.

In conclusion, we found that the transient hypertrophy induced by UDD is dependent on muscle fiber length, with longitudinal hypertrophy reducing the trigger for hypertrophy. The hypertrophy coincides with increased phosphorylation of the N2A element. The N2A-associated MARP proteins are strongly upregulated following UDD and deletion of the MARPs increases the extent of longitudinal hypertrophy following UDD, indicating the MARPs serve roles in inhibiting longitudinal hypertrophy. MARP tKO mice show altered regulation in the mTORC1 pathway following UDD and inhibition of mTORC1 by rapamycin shows mTOR is a main regulator of longitudinal hypertrophy.

## METHODS

### Animal studies

All experiments were done in accordance with the University of Arizona Institutional Animal Care and Use Committee and followed the US National Institutes of Health Using Animals in Intramural Research guidelines for animal use. We used 3-month-old C57BL/6J mice referred to as wildtype (WT), and 6-month-old Sprague Dawley rats (SD). Knockout models for titin binding proteins MARP1-3 (Ankrd1, Ankrd2 and Ankrd23)^10,44,53^ were kindly provided by Dr Ju Chen and Dr Stephan Lange. Homozygous Rbm20^ΔRRM^ mice ^21,54^ and Rbm20 ko rats^55,56^, have previously been described. Mice were maintained on a C57BL/6J background, with the data from the MARP mice being on a black swiss background.

#### Surgical procedure

For unilateral diaphragm surgery (UDD)^21^ or bilateral diaphragm denervation (BDD) studies, mice or rats were anaesthetized with 2-3% isoflurane and a small incision was made in the neck area just above the clavicle. The right phrenic nerve was isolated behind the sternohyoid muscle, and a 3–4 mm section was transected at the height of the supraclavicular nerve branch. For BDD surgery both left, and right phrenic nerves being transected. Sham operated animals underwent the same procedure, except the phrenic nerve was left intact. Animals were sacrificed 1, 3, 6, 12 or 35-days after surgery for morphometric analysis and tissue harvest.

### Pharmacological inhibition of hypertrophy

Inhibitor studies with rapamycin (mTOR inhibitor) and cyclosporin A (Calcineurin inhibitor) were performed by injecting mice, twice daily intra-peritoneal (IP), with 2.5 or 25 mg/kg/day, respectively. Each inhibitor was dissolved in Dimethylsulfoxide (DMSO) and diluted to 20% DMSO with saline solution just before injection. Mice received inhibitors or vehicle (20% DMSO in saline) starting 3-days prior to UDD surgery to prime the mice and continued following UDD to inhibit hypertrophy growth (FIG. 4C). Mice were sacrificed after 3-days UDD and processed for serial sarcomere measurements as described below.

### Serial sarcomere measurements

Previously described in van der Pijl et al^21^. Briefly, mice were anaesthetized with a 140/10 mg/kg ketamine/xylazine solution, a small incision to visualize the jugular vein, which was subsequently cannulated for perfusion. Mice were perfused with a solution consisting of 4% formaldehyde, with 70 U/mL of heparin in phosphate buffered saline (PBS), after which the diaphragm was removed and stored in 4% formaldehyde in PBS overnight for complete fixation. Full length diaphragm midcostal strips were gently dissected and flattened between glass slides, costal width was measured using a caliper and sarcomere lengths were measured using a He/Ne laser diffraction system.

Alternatively, muscle fiber bundels from chemically demembranated full length diaphragm midcostal strips of 3-day mice were flattened between glass slides. Costal width and sarcomere lengths were measured using a Zeiss Axio Imager M1 microscope (Zeiss), at ×50 magnification to measure the length of the muscle bundles and ×640 for sarcomere length along four points of the fiber bundles. Images were captured using AxioCam MRc with Axiovision software (Zeiss) and images were calibrated using a 0.01 mm stage micrometre (Edmund Optics). To determine the number of serial sarcomeres the muscle bundle length (costal width) was divided by the sarcomere length.

Demembranating solution consisted of a relaxing solution (in mM; 20 BES, 10 EGTA, 6.56 MgCl2, 5.88 NaATP, 1 DTT, 46.35 K-propionate, 15 creatine phosphate, pH 7.0), with 1% Triton-X-100 at 4°C, and protease inhibitors (phenylmethylsulfonyl fluoride (PMSF), 0.25 mM; leupeptin, 0.04 mM; E64, 0.01 mM, ), after demembranation, samples were stored in just relaxing solution plus inhibitors (without triton X-100) at 4°C.

### Transcriptome studies

RNA sequencing (RNAseq) was performed on right costal diaphragm samples collected from 3-day sham and UDD animals and flash frozen in liquid nitrogen until further processing. For RNA extraction, samples were incubated overnight in RNAlater-ICE (Thermo Scientific) and subsequently transferred to RLT buffer for extraction according to the RNeasy Fibrous Tissue Mini Kit (Qiagen). Tissue disruption was achieved using a Bullet Blender (Next Advance) and Green Eppendorf lysis kit tubes (Next Advance), by grinding samples for 4 minutes at setting 10. Thereafter, total RNA extraction was performed following the RNeasy Fibrous Tissue Mini Kit’s instructions and quantified using a Nanodrop ND-1000 spectrophotometer (Thermo Scientific). Each sample consisted of 3 biological replicate sham or UDD samples. Both Library preparation and sequencing was performed by the University of Chicago Genomics Facility, Chicago, USA. Briefly, library preparation: rRNA was depleted from RNA preparations from 1 µg total RNA. Libraries were prepared using an RNA Library Prep Kit from Illumina following the manufacturer’s instructions. Sequencing was performed on an Illumina Hiseq2500 sequencer using 100 bp paired-end sequencing. For RNAseq analysis see^20^. Briefly, Adapters and low-quality reads were removed with Trim Galore and reads were mapped to the rat genome (Release mRatBN7.2) using STAR^57^ with default settings. Differentially expressed genes were determined with DESeq2^58^. Genes with population adjusted p-values (p_adj_) <0.05 were considered differentially expressed. For titin splicing, percent spliced in index (PSI) was calculated as a measure for determining if an exon is spliced in, following the titin exon annotation by Bang et al.^59^. RNA sequence data were deposited at NIH NCBI BioProject (accession number PRJNA1460879).

### Preparation of muscle for mass spectrometry analysis

Diaphragm muscle was ground to a fine powder using Dounce homogenizers cooled in liquid nitrogen and acclimated to –20°C for 30 min before continuing. Tissue powder was resuspended at a concentration of 50 mg/ml in a Urea buffer (4 M urea, 1 M thiourea, 25 mM Tris–HCl, 75 mM dithiothreitol, 1.5 % SDS, 25 % glycerol, pH 6.8) with protease inhibitors (0.04 mM E-64, 0.16 mM leupeptin, and 0.2 mM PMSF). The solution was mixed for 4 min, followed by 10 min of incubation at 60°C. Samples were centrifuged at 12.000 rpm and the supernatant flash frozen for storage at −80°C.

### In-solution Tryptic Digestion

50 µg of rat costal diaphragm lysate was subjected to acetone precipitation by adding six times the sample volume of pre-chilled 100 % acetone and incubated one hour at -20°C. The precipitates were centrifuged at 16,000 x g for 10 minutes at 4°C and the acetone was removed.

400 µL of pre-chilled 90% acetone was added to the protein pellet, briefly vortexed and centrifuged at 16,000 x g for 5 minutes at 4°C. The remaining acetone was removed, the protein pellets were air dried for 3 minutes, resuspended in 100 µL of 50 mM NH_4_HCO_3_ and sonicated for 5 minutes. The samples were supplemented with dithiothreitol (DTT) at a final concentration of 5 mM and incubated at 70°C for 30 minutes. Samples were cooled to room temperature for 10 minutes and incubated with 15 mM acrylamide for 30 minutes at room temperature while protected from light. The reaction was quenched with DTT with a final concentration of 5 mM and incubated in the dark for 15 minutes. One µg of Lys-C was added to each sample and incubated at 37° C for 2-3 hours while shaking at 300 rpm followed by the addition of 50 µL of 50 mM ammonium bicarbonate and 2 µg of trypsin and incubation overnight at 37°C while shaking at 300 rpm. 14.7 µL of 40 % FA/1 % HFBA was added to each sample and incubated for 10 minutes (final concentration is 4 % FA/0.1 % HFBA) to stop trypsin digestion. The samples were desalted with Pierce Peptide Desalting Spin Columns per the manufacturer’s protocol (ThermoFisher Scientific, cat no. 89852) and the peptides were dried by vacuum centrifugation. The dried peptides were resuspended in 20 µL of 0.1 % FA (v/v) and the peptide concentration was determined with the Pierce Quantitative Colorimetric Peptide Assay Kit per the manufacturer’s protocol (ThermoFisher Scientific, cat no. 23275). 350 ng of the final sample was analyzed by mass spectrometry.

### Phosphoproteomics

To determine global differences in protein phosphorylation abundance between sham or UDD, 1 mL of protein lysate corresponding to 50 mg diaphragm per sample (pooled costal diaphragm of 2 mice) was subjected to in-solution tryptic digestion and phosphopeptide enrichment using sequential enrichment from metal oxide affinity chromatography per manufacturer’s protocol (Thermo Scientific, cat no. A32993 & A32992) similar to as previously described^60,61^. The dried peptides were resuspended in 20 µL of 0.1 % FA (v/v) and the peptide concentration was determined with the Pierce Quantitative Colorimetric Peptide Assay Kit per the manufacturer’s protocol. 350 ng of the final sample was then analyzed by mass spectrometry.

### Mass Spectrometry

HPLC-ESI-MS/MS was performed in positive ion mode on a Thermo Scientific Orbitrap Fusion Lumos tribrid mass spectrometer fitted with an EASY-Spray Source (Thermo Scientific, San Jose, CA). NanoLC was performed using a Thermo Scientific UltiMate 3000 RSLCnano System with an EASY Spray C18 LC column (Thermo Scientific, 50cm x 75 μm inner diameter, packed with PepMap RSLC C18 material, 2 µm, cat. # ES803); loading phase for 15 min at 0.300 µL/min; mobile phase, linear gradient of 1–34 % Buffer B in 119 min at 0.220 JL /min, followed by a step to 95% Buffer B over 4 min at 0.220 µL/min, hold 5 min at 0.250 µL/min, and then a step to 1 % Buffer B over 5 min at 0.250 µL/min and a final hold for 10 min (total run 159 min); Buffer A = 0.1 % FA/H_2_O; Buffer B = 0.1 % FA in 80 % ACN. All solvents were liquid chromatography mass spectrometry grade. Spectra were acquired using XCalibur, version 2.3 (ThermoFisher Scientific). A “TopSpeed” data-dependent MS/MS analysis was performed (acquisition of a full scan spectrum followed by collision-induced dissociation mass spectra of the Top N most intense precursor ions within the 3 second cycle time). Dynamic exclusion was enabled with a repeat count of 1, a repeat duration of 30 seconds, an exclusion list size of 500, and an exclusion duration of 40 seconds.

### Label-free Quantitative Proteomics

Progenesis QI for proteomics software (version 2.4, Nonlinear Dynamics Ltd., Newcastle upon Tyne, UK) was used to perform ion-intensity based label-free quantification as previously described^62^. In brief, in an automated format, raw files were imported and converted into two-dimensional maps (y-axis = time, x-axis =m/z) followed by selection of a reference run for alignment purposes. An aggregate data set containing all peak information from all samples was created from the aligned runs, which was then further narrowed down by selecting only +2, +3, and +4 charged ions for further analysis. The samples were then grouped according to treatment. Peak lists of the top ten fragment ion spectra were exported in Mascot generic file (mgf) format and searched against either the 2020_06 Swiss-Prot *Rattus norvegicus* database (8128 entries) , the 2018_11 Swiss-Prot *Mus musculus* database (17008 entries), or species respective TrEMBL databases using Mascot (Matrix Science, London, UK; version 2.6.0). The search variables that were used were: 10 ppm mass tolerance for precursor ion masses and 0.5 Da for product ion masses; digestion with trypsin; a maximum of two missed tryptic cleavages; variable modifications of oxidation of methionine, phosphorylation of serine, threonine, and tyrosine, and carbamidomethylation of cysteine; 13C = 1. The resulting Mascot .xml file was then imported into Progenesis, allowing for peptide/protein assignment, while peptides with a Mascot Ion Score of <25 were not considered for further analysis. False discovery rate was set to 1% for both peptides (minimum length of 7 amino acids) and proteins. Abundances were normalized to the total ion current (TIC) to correct for differences in sample loading and instrument response. For quantification, proteins must have possessed at least one or more unique, identifying peptide.

### Visualization and analysis of transcriptome and proteome data

Figures were generated using online resources, briefly, quantitative Venn diagrams were made using https://www.biovenn.nl/^63^. Lolipop graphs of gene onthology (GO) enritchment analysis by ShinyGO v0.75 http://bioinformatics.sdstate.edu/go/^64^. Heatmaps were generated using a combination of Graphpad Prism v9.1, Perseus v2.0.3.1^65^ and Heatmapper (http://heatmapper.ca/)^66^. Perseus v2.0.3.1 was also used to analyze principal components and visualized using Graphpad Prism v9.1. A protein mapping table was constructed in R, using the R Interface to UniProt Web Services package version 2.34.0.^67^ UniprotKB identifiers and gene ID’s were assigned to each peptide. Phosphorylated peptides were mapped to the UniProt sequences to identify the positions of the modified residue(s) using stringr version 1.4.0.^68^ For representation of titin phosphorylation, normalized abundance values were converted to Z-scores by subtracting the grand-mean of normalized abundance values for a particular peptide and dividing by the grand-mean standard deviation. We subsequently summed Z-scores of isobaric peptides for a given phospho-site to generate values to represent expression levels. To generate domain or regional phosphorylation we further summed phospho-sites to derive domain and regional phosphorylation graphs.

### Western blot

Western blot experiments previously described in van der Pijl et al^21^. Proteins were transferred onto Immobilon-P PVDF 0.45 μm membranes (Millipore) using semi dry transfer (Bio-Rad).

Membranes were blocked with Odyssey blocking buffer (Li-Cor Biosciences) for 1 hour, and subsequently probed with primary antibodies at 4°C overnight (See Table 1). Near Infra-Red dyes were used as secondary antibodies for detection with Odyssey CLx Imaging System (Li-Cor Biosciences, United states).

**Table 1.** Antibodies used in this study.

| Antibody | Source/isotype | Dilution | Company | Catalog# |
| --- | --- | --- | --- | --- |
| MAPK1/3 (ERK2/1) | Mouse IgG1 | 1:200 | Cell Signaling | #4696 |
| Calcineurin | Mouse IgG2a | 1:750 | BD biosciences | 610260 |
| mTOR | Rabbit IgG | 1:750 | Cell Signaling | #2983 |
| P70S6K | Rabbit IgG | 1:500 | Cell Signaling | #2708 |
| 4EBP1 | Rabbit IgG | 1:750 | Cell Signaling | #9452 |
| Gapdh | Rabbit IgG | 1:5000 | Cell Signaling | #2112 |
| Gapdh | Mouse IgG1 | 1:3000 | Thermo Fisher Sci. | MA5-15738 |
| CF790 Goat anti Mouse IgG | Goat IgG | 1:10.000 | Biotium | 20342 |
| CF680 Goat anti Mouse IgG | Goat IgG | 1:10.000 | Biotium | 20065 |
| CF680 Goat anti Rabbit IgG | Goat IgG | 1:10.000 | Biotium | 20067 |
| CF680R Goat anti Mouse IgG2a | Goat IgG | 1:10.000 | Biotium | 20842 |

---

**S FIGURE 1.**
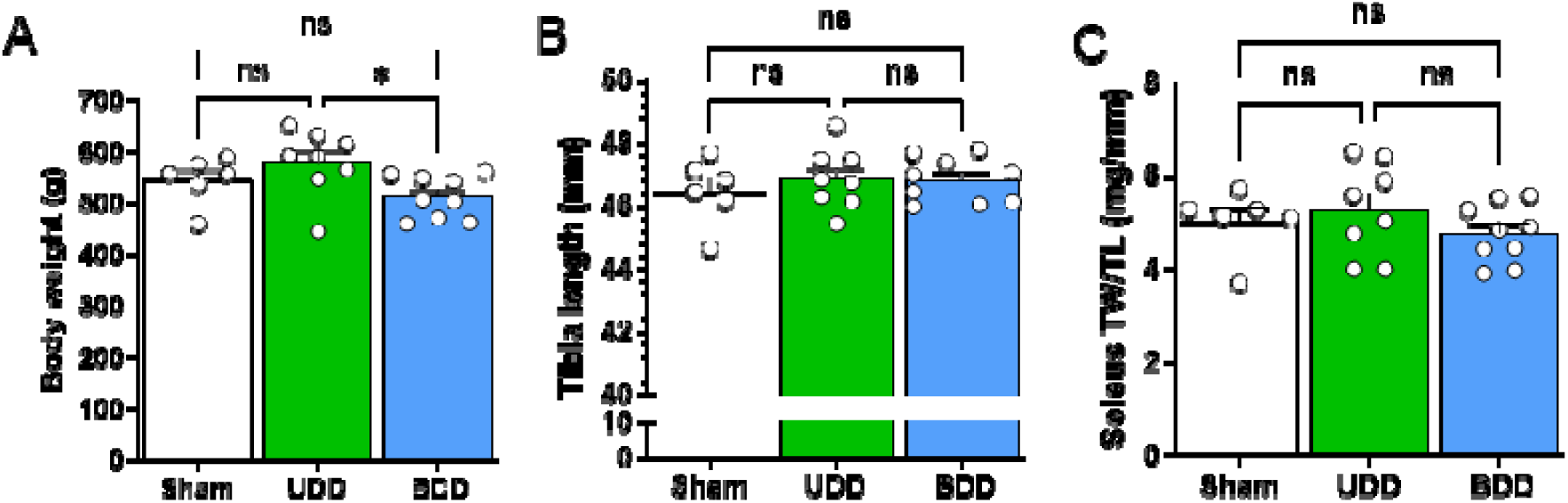
3-days bilateral diaphragm denervation in rats showed similar body weights compared to sham animals (A; n=6-9/group) and were of similar size based on tibia length (B) and soleus muscle weights (C). Statistical testing by one-way ANOVA and Dunnett’s multiple comparisons test.

**S FIGURE 2.**
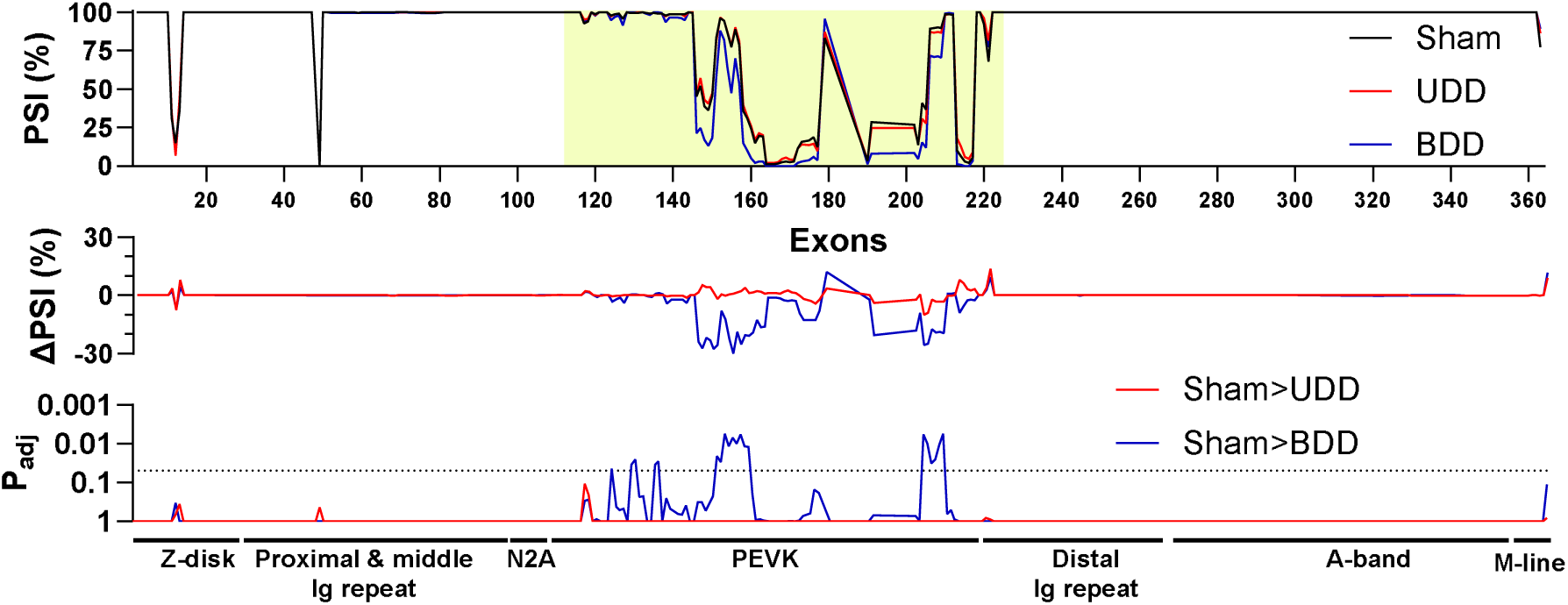
Titin splicing in 3-day unilateral and bilateral diaphragm denervation in rats showed similar levels of splicing between UDD (red line) and sham (black line) animals, while BDD (blue line) animals showed a decrease in exons coding for the elastic PEVK element. Statistical testing by Multiple t-testing, comparing sham to UDD and sham to BDD (n=4/group).

**S FIGURE 3.**
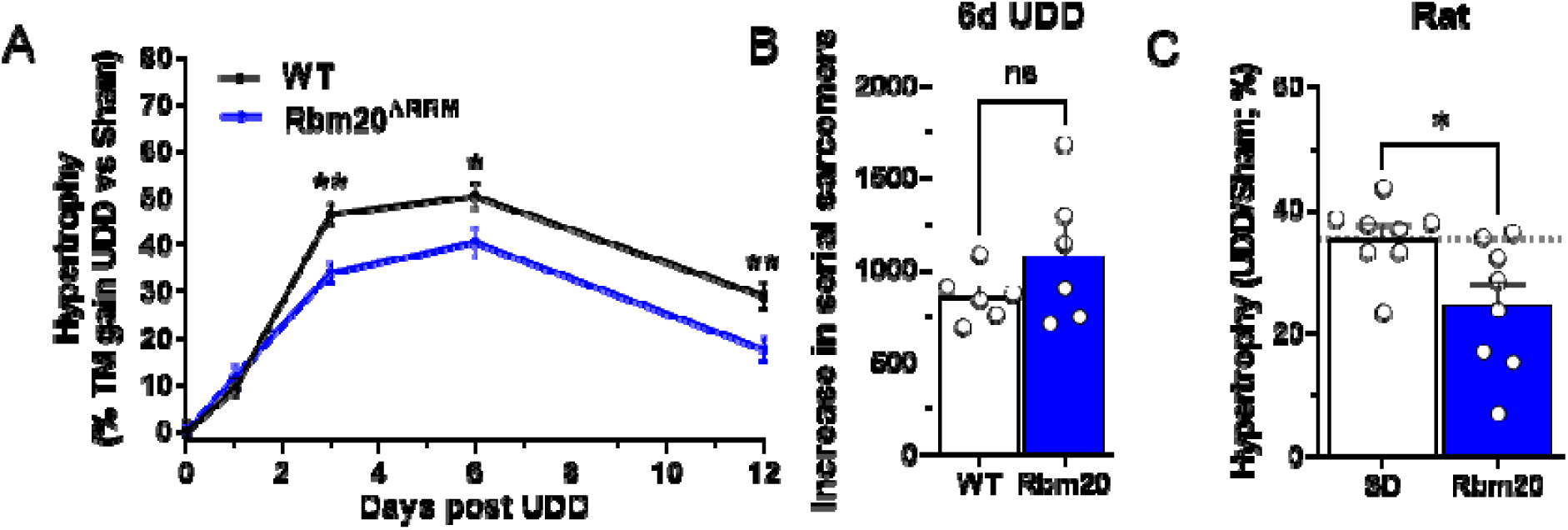
Role of titin stiffness on hypertrophy following UDD. (A) Transient hypertrophy response in Rbm20^ΔRRM^ mice (more compliant titin) showing a blunted hypertrophy response compared to WT mice, based on percent increase of diaphragm right costal mass relative to sham (n=10-12). (B) Titin-based stiffness does not alter longitudinal hypertrophy response, as both WT and Rbm20^ΔRRM^ mice show a similar increase in serial sarcomeres following 6-days UDD. Rbm20-KO rat response to 3-days UDD, based on percent increase of diaphragm right costal mass relative to sham (mouse n=10-11, rat n=8) supporting titin-based stiffness regulating muscle hypertrophy similarly across species. Statistical testing by t-test.

**S FIGURE 4.**
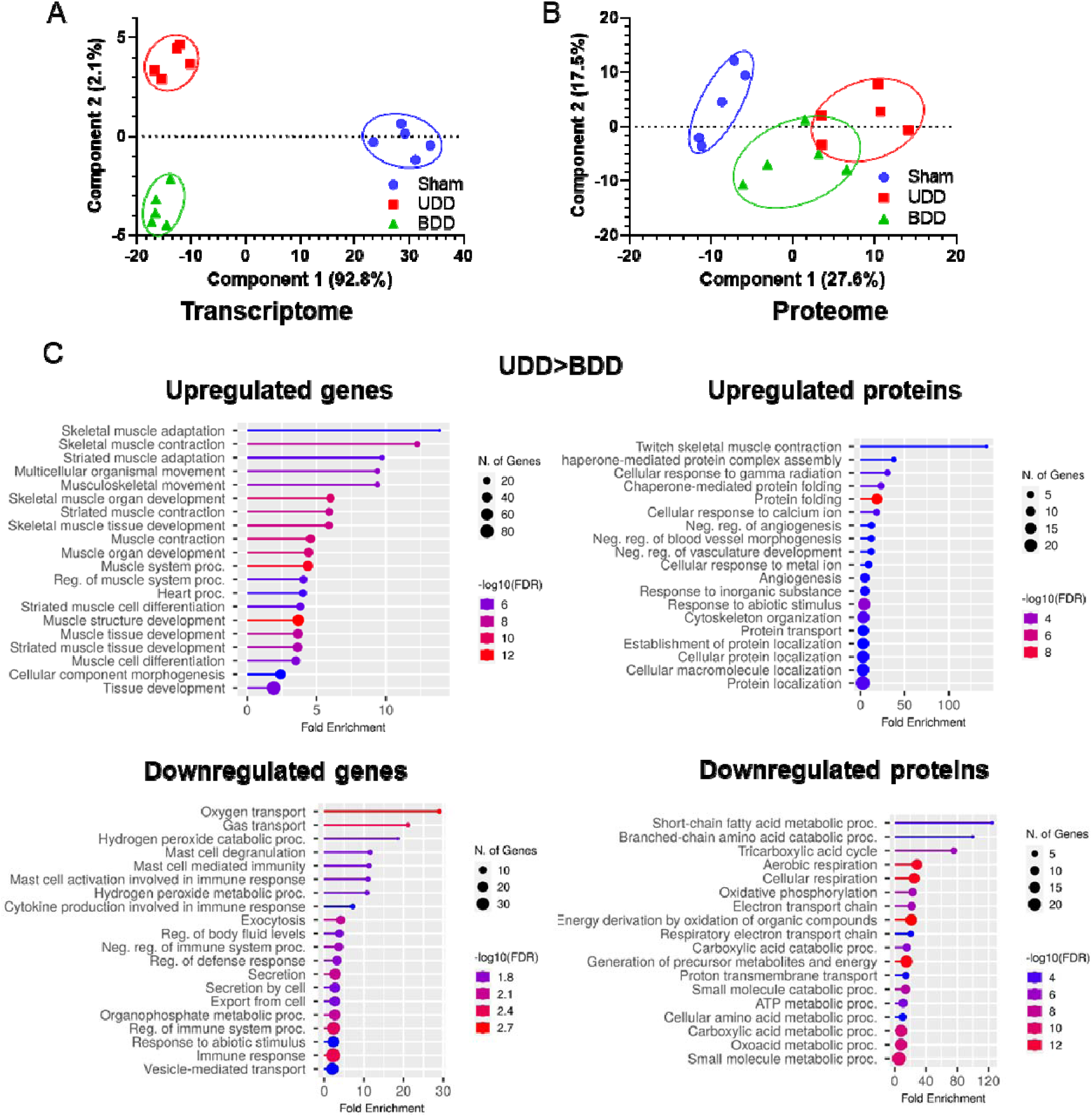
Principal component analysis of the rat 3-day UDD and BDD transcriptome (A) and proteome (B), showing clear separation of groups at the transcript level and overlap of BDD and UDD samples at the protein level. GOterm enrichment of UDD>BDD separated by up- or down-regulated transcriptomes and proteome (C, left and right, respectively) show distinct, yet overlapping cellular processes. Global mass spectrometry was analyzed by ANOVA and corrected for multiple comparisons with false discovery rate with a cut-off at p<0.05.

**S FIGURE 5.**
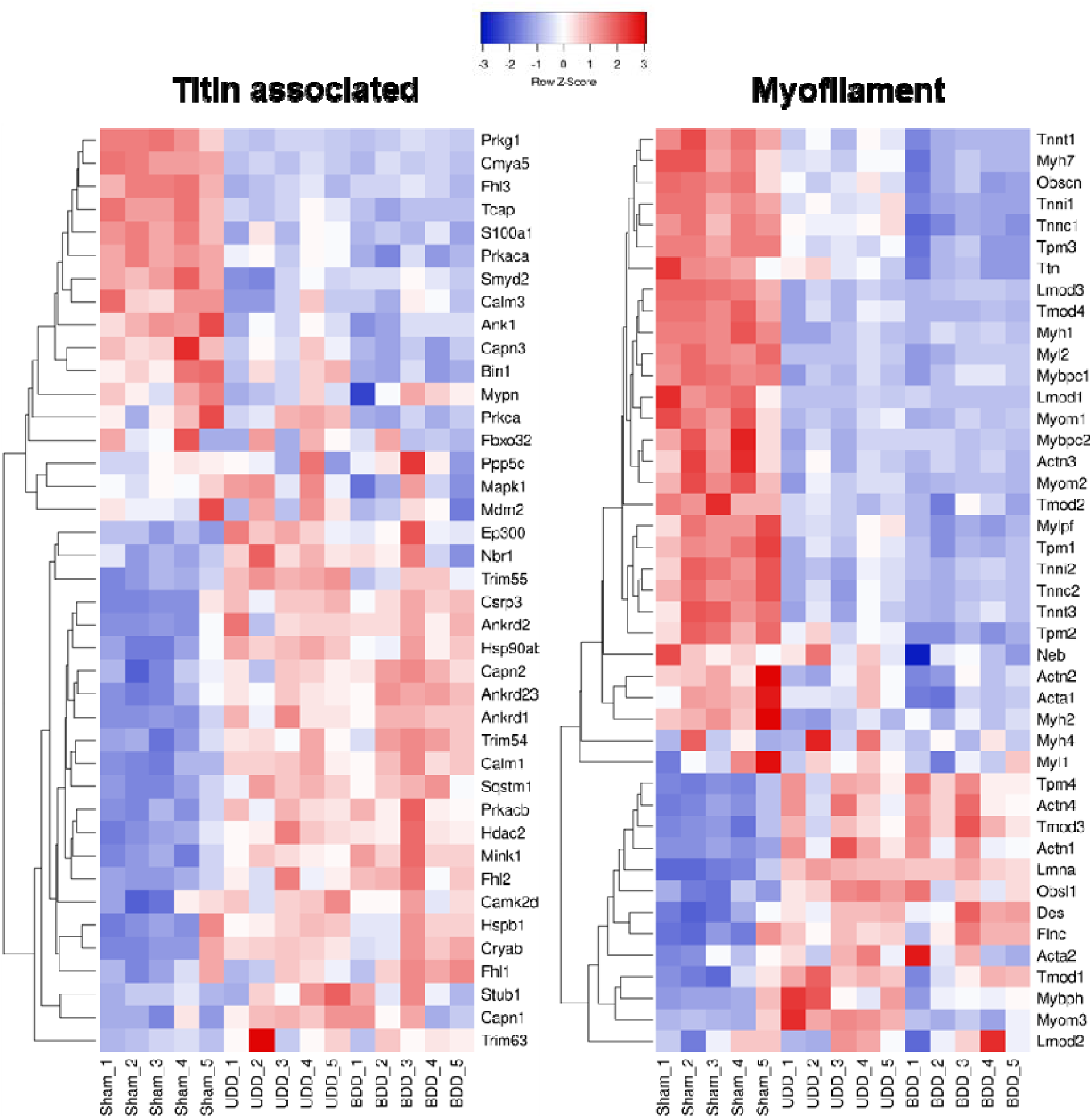
Transcriptome regulation of titin-associated and myofilament genes by RNAseq in rats following 3-days of UDD/BDD. Heatmaps showing similar regulation between UDD and BDD samples (n=4-5; Z-score: red= upregulated, blue= downregulated) at the transcript level for titin-associated and myofilament genes, based on hierarchal clustering.

**S FIGURE 6.**
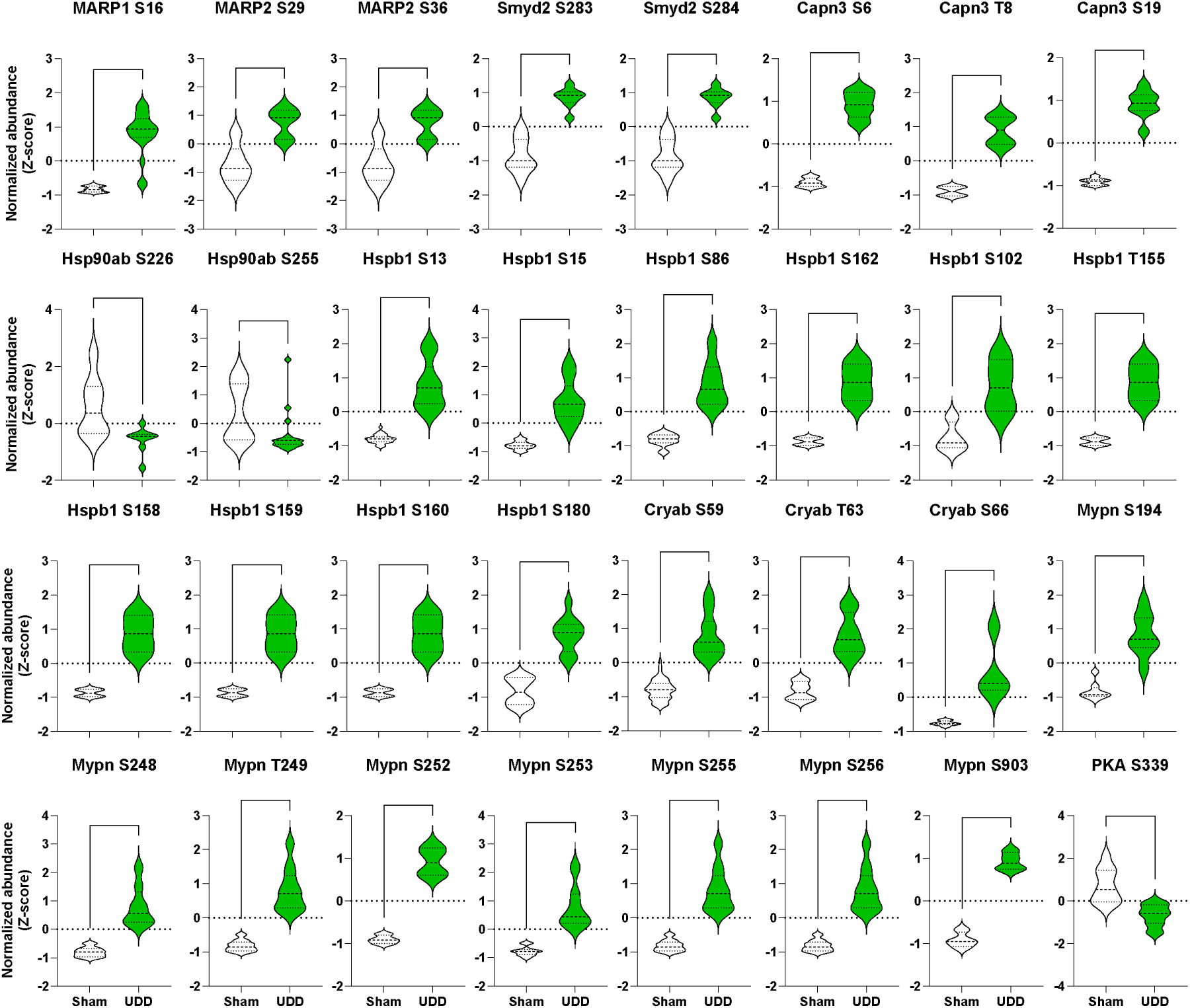
Titin N2A associated protein phosphorylation events at 24-hour UDD. Violin plots of phosphorylation events in N2A-associated proteins following UDD: MARP1 (Transcript: ENSMUST00000237142.2 Ankrd1-205), MARP2 (Transcript: ENSMUST00000026172.3 Ankrd2-201), Smyd2 (Transcript: ENSMUST00000027897.8 Smyd2-201), Capn3 (Transcript: ENSMUST00000028749.15 Capn3-202), Hsp90ab (Transcript: ENSMUST00000024739.14 Hsp90ab1-201), Mypn (Transcript: ENSMUST00000095580.3 Mypn-201), Hspb1 (Transcript: ENSMUST00000005077.7 Hspb1-201), Cryab (Transcript: ENSMUST00000217475.2 Cryab-206) and Prkca/PKA (Transcript: ENSMUST00000005606.8 Prkaca-201). Data represented as Log2 of the normalized abundance with significance determined by Kolmogorov-Smirnov test.

**S FIGURE 7.**
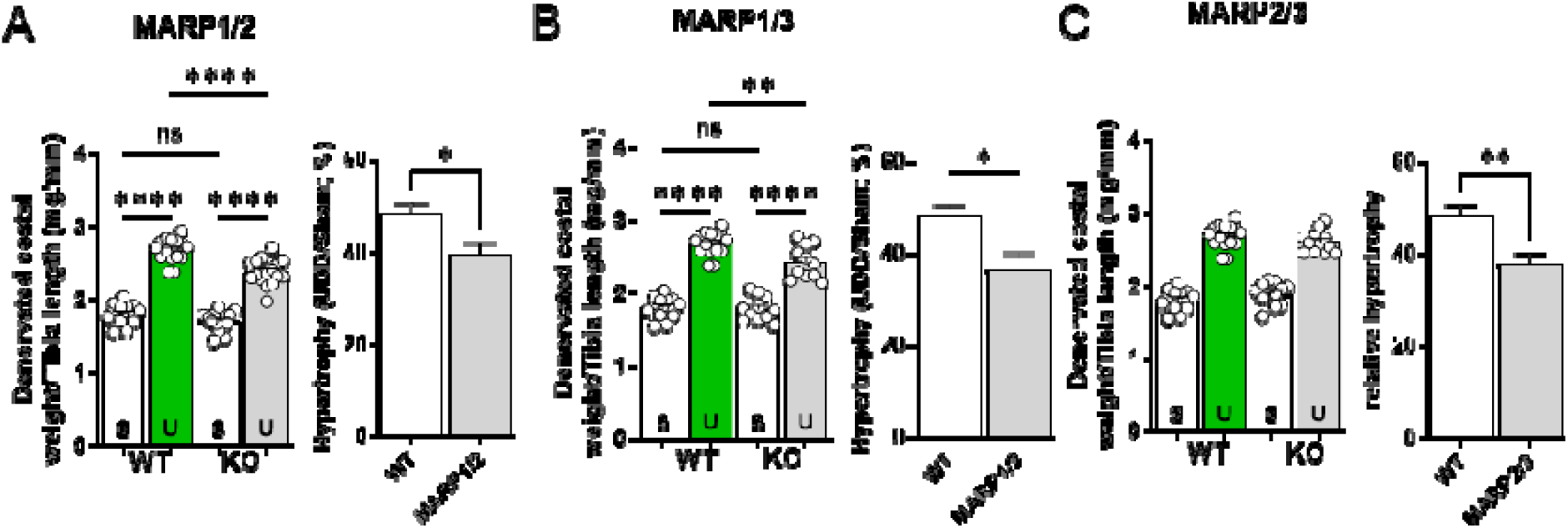
6-day UDD on double KO mice of MARPs. Double KO of MARP1/2 (A), MARP1/3 (B) and MARP2/3 (C) all showed a reduction in hypertrophy following UDD, suggesting redundancy between the MARPs. Left panel, diaphragm right costal mass normalized to tibial length and right panel, percentual increase in right costal mass relative to sham. S= Sham, U= UDD (n=10-12).

